# Maturation of the matrix and viral membrane of HIV-1

**DOI:** 10.1101/2020.09.23.309542

**Authors:** Kun Qu, Zunlong Ke, Vojtech Zila, Maria Anders-Össwein, Bärbel Glass, Barbara Müller, Hans-Georg Kräusslich, John A. G. Briggs

**Author notes:** Equal contributions (Zunlong Ke, Vojtech Zila).

## Abstract

Gag – the main structural protein of HIV-1 – is recruited to the plasma membrane for virus assembly by its matrix (MA) domain. Gag is subsequently cleaved into its component domains, causing structural maturation to repurpose the virion for cell entry. We determined the structure and arrangement of MA within immature and mature HIV-1, providing a basis to understand MA’s role in virus assembly. Unexpectedly, we found that MA rearranges during maturation, to form a new, hexameric lattice in which the acyl chain of a phospholipid extends out of the membrane to bind a pocket in MA. Our data suggest that proteolytic maturation of HIV-1 not only achieves assembly of the viral capsid surrounding the genome, but extends to repurpose the membrane-bound MA lattice for an entry or post-entry function, and causes partial removal of 2,500 acyl chains from the viral membrane.

## Introduction

Assembly and budding of HIV-1 is initiated at the plasma membrane (PM) and driven primarily by a 55-kDa viral polyprotein named Gag. Gag consists of an N-terminal matrix (MA) domain, responsible for recruitment to the PM, the capsid (CA) domain, which induces Gag self-assembly *via* protein-protein interactions, the nucleocapsid (NC) domain, which recruits the viral RNA genome to the assembly site, as well as some minor peptide domains (*1, 2*). Protein-protein, protein-lipid and protein-RNA interactions lead to clustering of Gag at the assembly site, membrane bending, and subsequent release of the membrane-enveloped immature HIV-1 particle. Concomitant with, or shortly after release, the viral protease (PR) cleaves Gag at multiple positions leading to a dramatic structural rearrangement to repurpose the virus particle for entry into a target cell. Maturation results in condensation of a ribonucleoprotein complex (RNP) from NC and RNA, which becomes surrounded by the cone-shaped capsid made of CA, while mature MA is thought to remain associated with the viral membrane. Rearrangement of the HIV-1 envelope (Env) glycoprotein on the particle surface is presumed to render the virion fusogenic(*2, 3*).

The structure of the heterologously-expressed, 17-kDa MA protein has been determined, revealing a small folded domain composed of five alpha helices and one 3_10_ helix between helices 2 and 3 (*4, 5*). MA crystallizes as a trimer (*5*), and multimerizes on artificial membrane monolayers into a hexameric lattice of trimers with holes or gaps at the six-fold symmetric positions in the lattice (*6*). These holes have been suggested to provide binding sites for the C-terminal tail of HIV-1 Env and promote Env incorporation into assembling virions (*7, 8*).

Membrane recruitment of Gag *via* MA is mediated by an N-terminal myristate moiety as well as by a highly basic region (HBR, residues 17-31) (*9, 10*). The N-terminal myristate is thought to be in equilibrium between sequestration in a pocket in the MA domain, and an exposed conformation (*11*) – the so-called myristoyl switch. Myristate exposure can be promoted by trimerization of MA (*11*). Additionally, binding of PI(4,5)*P_2_* in an extended-lipid conformation to a pocket on the side of MA had been proposed to promote exposure of the myristate and its insertion into the lipid bilayer (*12*). This mode of PI(4,5)*P_2_* binding, in which the 2’ acyl chain of the lipid would have to be pulled out of the bilayer, is no longer widely considered relevant to MA-membrane interactions during virus assembly, which instead are likely to be between the PI(4,5)*P_2_* headgroup and the membrane-facing basic MA residues (*13–15*). MA interacts with nucleic acids in the cytosol, in particular tRNA, *via* the HBR (*16, 17*). These observations support a model in which exchange of nucleic acid for the PI(4,5)*P_2_* head group promotes myristate exposure and stabilization of PM binding during virus assembly (*16, 17*).

The known roles of MA are performed during virus assembly, and no function for MA during entry or post-entry stages is currently established. Models for how MA functions during assembly are limited by the current lack of structural information on MA within virions. It is unclear what conformation and arrangement MA adopts *in situ*, and whether MA undergoes any structural changes in the process of virus maturation. Here, we set out to determine the structure and arrangement of MA within immature and mature virus particles.

## Results and Discussion

Purified mature particles generated by transfection of a non-infectious HIV-1 proviral derivative that expresses all viral proteins except for Nef (pcHIV) (*18*), and immature particles generated by transfection of pcHIV carrying an active site mutation in PR (pcHIV PR-) were imaged without chemical fixation using cryo-electron tomography (cryo-ET). Visual inspection of tomograms revealed the expected features: a striated CA layer in immature virus particles, and conical CA cores in mature virus particles (**Fig. 1A-B**). In some regions underlying the lipid envelope of mature virions, we observed regular lattice-like features (**Fig. 1C**).

**Fig. 1.**
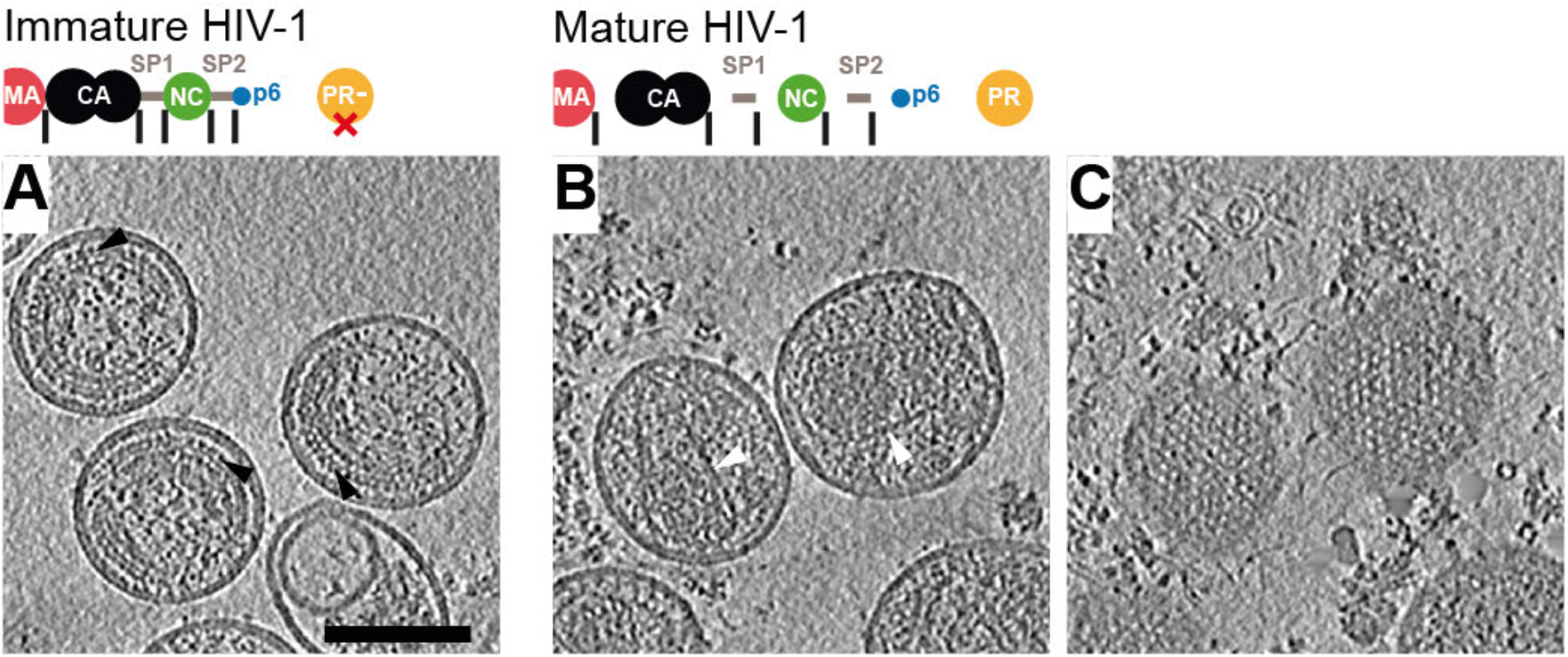
Tomograms of immature and mature HIV-1 particles. (**A**) Top: Schematic of the Gag protein domain architecture with cleavage sites marked as vertical black lines. The D25N mutation (red X) inactivates the viral PR. Bottom: Computational slices through a representative tomogram of immature HIV-1 (cHIV PR-) particles. Density is black. The striated Gag layer is distinctly observed in immature particles (black arrowheads). **(B)** As in (A) for cleaved, mature HIV-1 (cHIV) particles. Distinct conical CA cores are observed (white arrowheads). (**C**) slices grazing the inner surface of the membrane at the top of the particles in (B) revealing a regular MA lattice. Gold fiducials are removed from the images. Scale bar: 100 nm.

We subjected the region underlying the viral membrane in immature cHIV particles to reference-free subtomogram averaging. This analysis revealed a hexameric lattice of MA trimers with a hexamer-hexamer spacing of 9.8 nm containing large holes at the six-fold axes (**Fig. 2A**), reminiscent of the lattices observed for membrane-associated purified MA in vitro (*6*). In order to improve the spatial resolution of the immature MA structure, we analysed a cryo-ET dataset of fixed, immature, complete HIV-1_NL4-3_ particles (*19*). From this dataset, we obtained the same MA lattice structure at 7.2 Å resolution (**Fig. 2B-F, Fig. S1**). The poorly-ordered and sparse MA lattice covered only small patches of the inner membrane leaflet, while in other areas MA did not appear to form a lattice, suggesting heterogeneity in MA packing (**Fig. 2G, Fig. S2**). Despite this heterogeneity, we could clearly resolve alpha-helices (**Fig. 2C-F**). The MA trimer structure determined by crystallography (PDBID: 1HIW) (*5*) could be fitted into the density as a rigid body. The MA domain of Gag is therefore present as a trimer within immature HIV-1 particles (**Fig. 2D, Fig. S3A**).

**Fig. 2.**
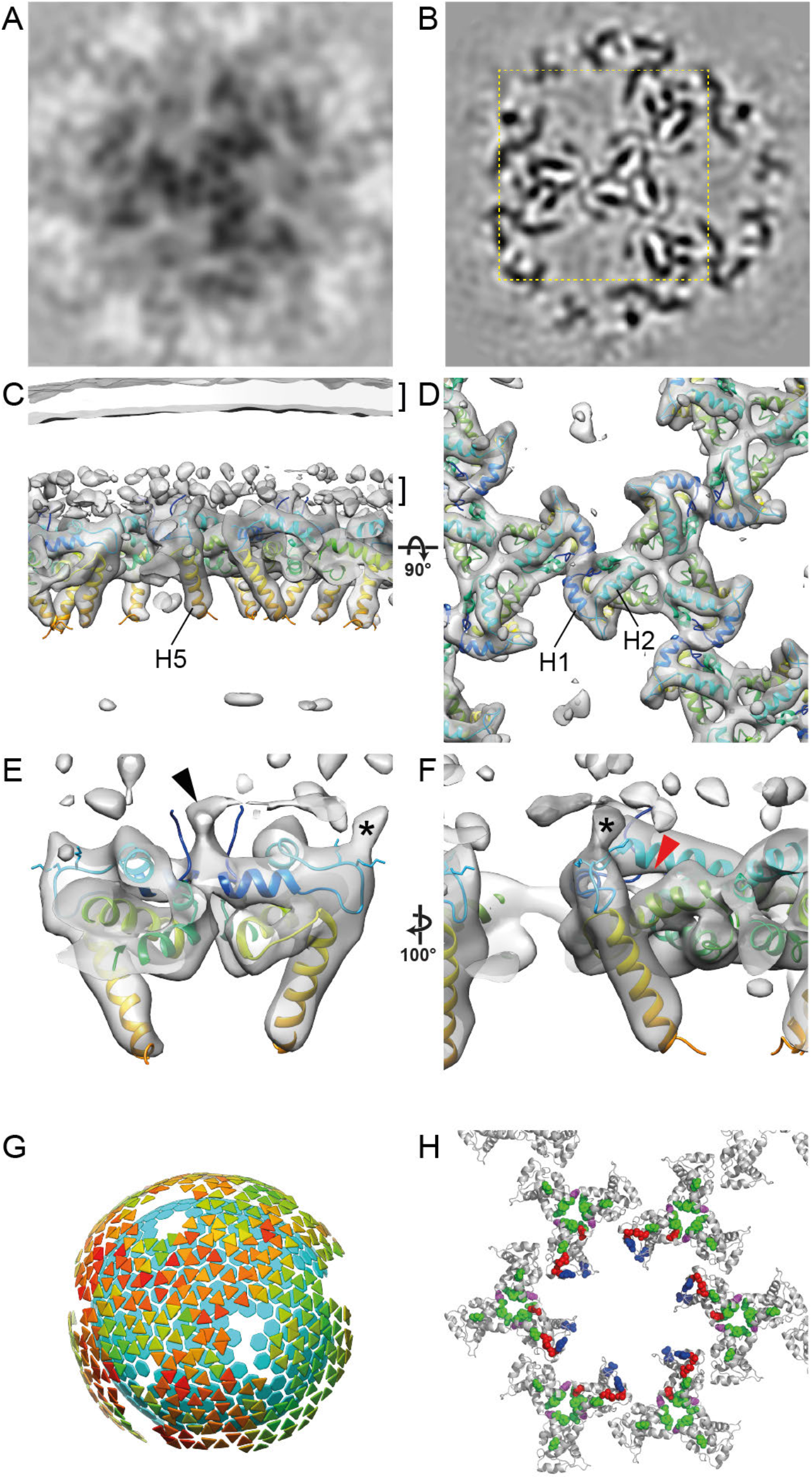
The immature HIV-1 matrix structure. **(A, B)** Slices through reconstructions of the immature HIV-1 MA lattice in cHIV PR- (A) and HIV-1_NL4-3_ PR- (B). The boxed region in B is shown at a higher magnification in (D). Density is black. **(C, D)** Isosurface views of the cryo-ET reconstruction (grey) for immature HIV-1_NL4-3_ PR-MA lattice cut perpendicular to the membrane (C) or viewed from the top towards the membrane center (D). The two layers of density corresponding to the lipid headgroup layers are indicated by brackets. The structure of monomeric MA determined by NMR (PDBID: 2H3Q; colored blue to red from N to C terminus) was fitted as a rigid body into the density. Helices 1, 2 and 5 are marked. **(E)** As in (C), enlarged and cut to reveal density corresponding to the N-terminal residues (black arrowhead). The density marked with an asterisk corresponds to K25 and/or K26. **(F)** As in (E) but rotated. The red arrowhead indicates the unoccupied PI(4,5)*P_2_* binding pocket. **(G)** Lattice map derived from subtomogram averaging for the immature HIV-1_NL4-3_ PR-MA lattice. Lattice maps illustrate positions and orientations of MA trimers as triangles colored on a scale from red (lower cross correlation to average structure) to green (higher cross correlation to average structure). The positions of CA hexamers are indicated as blue hexagons to illustrate the relationship between MA and CA layers. **(H)** Immature MA lattice is shown as grey ribbons and residues whose mutation has been reported to modulate Env incorporation are shown as colored spheres. Mutations at residues L12, L30 and L74 (red), E16 and E98 (blue), and T69 (purple) impair Env incorporation. Mutations in V34, F43, Q62 and S66 (green) can rescue Env incorporation defects, except those caused by mutations at T69 (purple). E16 and E98 (blue) face the hole in the MA lattice in the immature virion.

MA helices 1 and 2 lie approximately parallel to the membrane and form a surface rich in charged and hydrophobic residues by which MA is attached to the inner membrane leaflet (**Fig. 2C, F**). MA trimers are packed together to form a lattice by a novel dimeric interface where the N-terminal residues and the N-terminus of helix 1 interact with themselves and the 3_10_ helix (**Fig. 2D, E**). We did not observe any density corresponding to PI(4,5)*P_2_* in the described binding site on the side of MA next to the HBR (*12*) (**Fig. 2F**). Its absence is consistent with current models in which the head group of PI(4,5)*P_2_* functions as the major cellular determinant of efficient MA-PM targeting by binding to the membrane-proximal face of MA (*13, 20*). The N-terminal region (residues 1-11) that changes conformation during the myristoyl switch (*11, 21*), is directly involved in the trimer-trimer interactions (**Fig. S3B**). At lower isosurface thresholds, a connection to the membrane appears in the vicinity of the N-terminus of helix 1 (**Fig. 2E**), while no notable density was observed in the myristoyl pocket (**Fig. S3C**). The conformational equilibrium is therefore shifted towards the exposed, membrane-inserted conformation in the immature virion, consistent with the myristate moiety becoming exposed upon MA oligomerization or membrane recruitment.

The C-terminal helix 5 of MA extends towards the centre of the virus particle and is separated from the underlying CA layer by a disordered stretch containing the MA/CA cleavage site (**Fig. 2C**). CA is therefore not resolved when MA is aligned, and equivalently MA is not resolved when CA is aligned (**Fig. S4**). The position of MA is thus constrained but not fixed relative to CA.

A number of studies have implied that holes in a hypothetical MA lattice may present a binding site for the cytoplasmic tail of Env, and mutations affecting MA trimerization have been shown to affect Env incorporation (*6, 7, 22*). Here we found that MA does form a lattice containing holes. We did not observe density for the Env tail in the holes, but that is not to be expected given the low Env copy number per particle (*23*). A number of mutations in MA have been reported to result in defects in Env incorporation (*7, 8, 22, 24–27*) (**Fig. 2H**, red, purple and blue spheres). Many of the mutated residues (red and purple spheres) are not exposed towards the central hole, but are rather located at sites close to the N-terminal residues of MA or the intra- and inter-trimer interfaces consistent with their phenotype in MA trimerization (**Fig. 2H**). These mutations are thus likely to modulate MA trimerization or lattice formation, and/or the myristoyl switch, but would not be presumed to directly control Env cytoplasmic tail binding within the holes. This is consistent with the model that correct MA oligomerization is required for Env incorporation (*8, 28*).

We next subjected the region underlying the membrane in mature cHIV particles to reference-free subtomogram averaging. This analysis revealed large patches of a hexagonal lattice of MA trimers with a hexamer-hexamer spacing of ~8.8 nm. These dimensions are similar to the 9 nm repeating MA lattice observed in anomalous, multi-cored, membranous particles (*29*). We resolved the structure of the lattice to a resolution of 9.5 Å (**Fig. 3A**). The mature lattice, like the immature lattice, is formed from MA trimers arranged in a hexagonal lattice, though it has a higher degree of regularity than the immature MA lattice (**Fig. S2**). The arrangement of MA within the immature and mature MA lattices is, however, strikingly different (**compare Figs. 2 and 3**) – the MA lattice undergoes structural maturation to form a new, different hexagonal lattice in the mature virion.

**Fig. 3.**
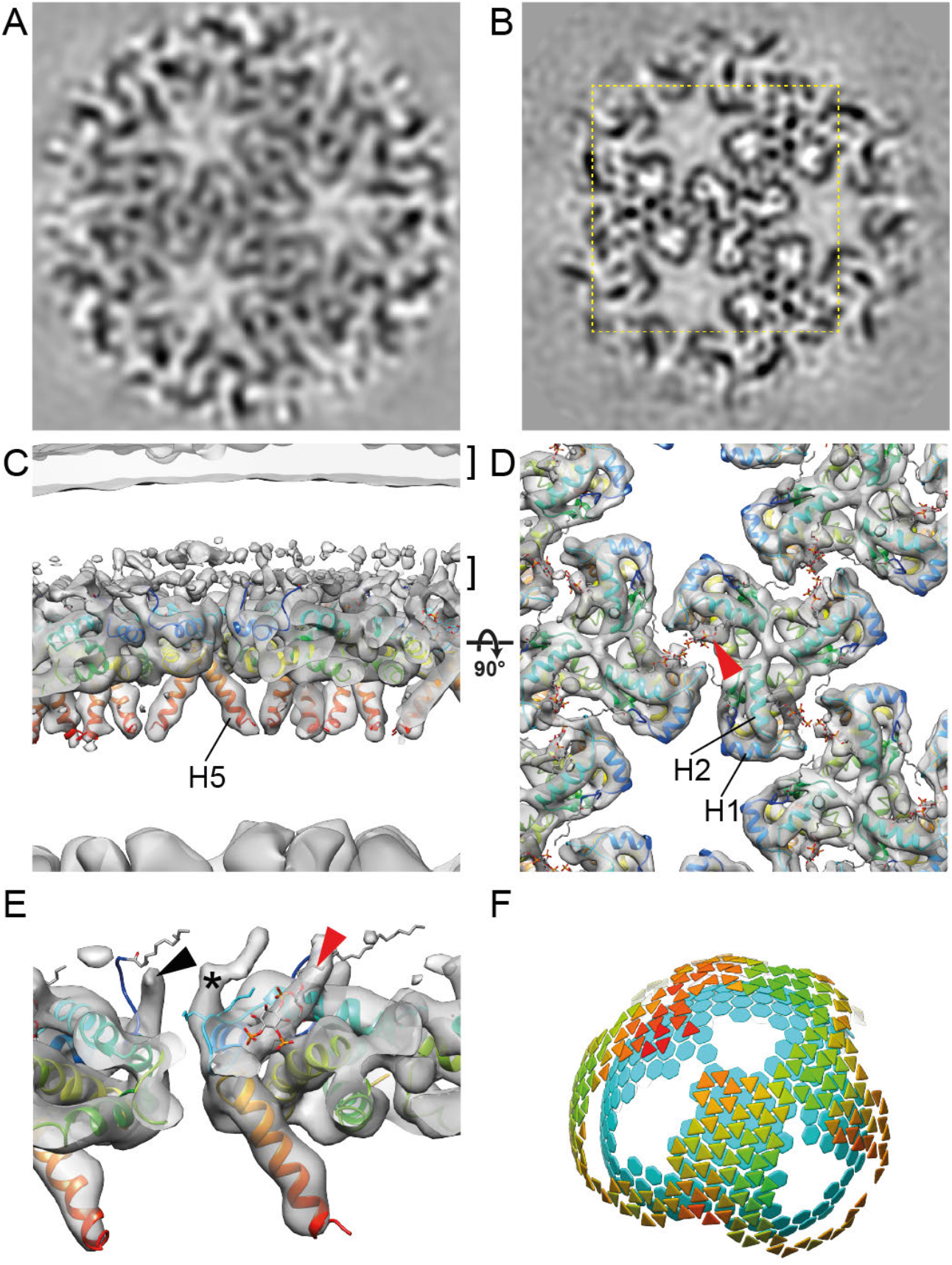
The mature HIV-1 matrix structure. **(A, B)** Slices through reconstructions of the mature HIV-1 MA lattice in cHIV (A) and cHIV MA-SP1 (B). The boxed region in B is shown at a higher magnification in (D). **(C, D)** An isosurface view of the cryo-ET reconstruction for mature cHIV MA-SP1 MA lattice fitted with the structure of monomeric MA, cut perpendicular to the membrane (C) or viewed from the top towards the membrane center (D). The two layers of density corresponding to the lipid headgroup layers are indicated by brackets. The structure of monomeric MA determined by NMR (PDBID: 2H3Q; colored blue to red from N to C terminus) was fitted as a rigid body into the density. Helices 1, 2 and 5 are marked. Density is observed in the PI(4,5)*P_2_* binding site (red arrowhead), and the structure of PI(4,5)*P_2_* as resolved bound to MA by NMR (PDBID: 2H3V), is shown as a stick model. **(E)** As in (C), enlarged and cut to reveal density corresponding to the N-terminal residues (black arrowhead), and PI(4,5)*P_2_* (red arrowhead). The density marked with an asterisk corresponds to K25 and/or K26. **(F)** As in Figure 2G, lattice map for the mature cHIV MA-SP1 MA lattice and the underlying CA lattice.

We had previously determined the structure of the CA layer within cHIV derivatives in which the PR cleavage site between MA and CA was blocked by mutation as potential maturation intermediates (HIV-1 MA-CA and MA-SP1) (*30*). CA and the CA lattice remain structurally immature in MA-CA and MA-SP1. We now analysed the MA layer in these datasets, and found that the MA lattice in HIV-1 MA-CA and MA-SP1 corresponds to that seen in mature particles, despite the absence of cleavage between MA and CA (**Fig. 3B, Fig. S5**). Cleavages downstream of SP1 are therefore sufficient for MA lattice maturation, which can occur without CA lattice maturation. It is unlikely that proteolytic cleavage between CA and NC could directly trigger MA maturation via a structural signal passed through CA without inducing any detectable structural change in CA. We therefore suggest that maturation of MA must be triggered “in trans” by another effector in the viroplasm or in the viral membrane. Further experiments will be required to elucidate the mechanism of MA maturation. Immature virions are fusion incompetent, and cleavages downstream of CA are sufficient to overcome this defect (*31*). These observations suggest that structural maturation of MA, rather than of CA, correlates with HIV-1 becoming fusogenic, potentially by allowing Env to redistribute on the virus surface(*32*).

We obtained a higher-resolution, 7.0 Å structure of the mature MA lattice from MA-SP1 particles, and - as for the immature MA lattice - the crystallographic trimer could be fitted into the density as a rigid body (**Fig. 3B-E, Fig. S1**). Comparison of the immature and mature MA structures revealed important differences (**Fig. 4**). In the immature lattice the HBR faces the holes at the six-fold axes, which are therefore surrounded by a basic surface (**Fig. 4B**). In contrast, in the mature virus, MA presents a largely neutral surface towards the holes (**Fig. 4B**). In the mature virus, basic residues in the HBR loop face acidic residues in the N-terminus of helix 4 (72E) and the 3_10_ helix (51E) of the adjacent MA monomer to form the dimeric interface that links trimers together (**Fig. 3D, Fig. 4A, Fig. S3B**). Unlike in the immature virus, where the lipid binding pocket on the side of MA is empty, in the mature virus it contains a density at the position previously described for PI(4,5)*P_2_*(PDBID: 2H3Q and 2H3V) (*12*). PI(4,5)*P_2_* can be modelled at this position in the extended-lipid conformation with the 2’ acyl chain removed from the bilayer, while the other acyl chain extends upwards towards the bilayer (**Fig. 3D, E, Fig. S6**). The PI(4,5)*P_2_* binding pocket is adjacent to the HBR, right at the centre of the dimer interface. The N-terminus of helix 1 prominently protrudes up into the PM, indicating that the myristate moiety retains an exposed, membrane-inserted conformation in the mature MA lattice.

**Fig. 4.**
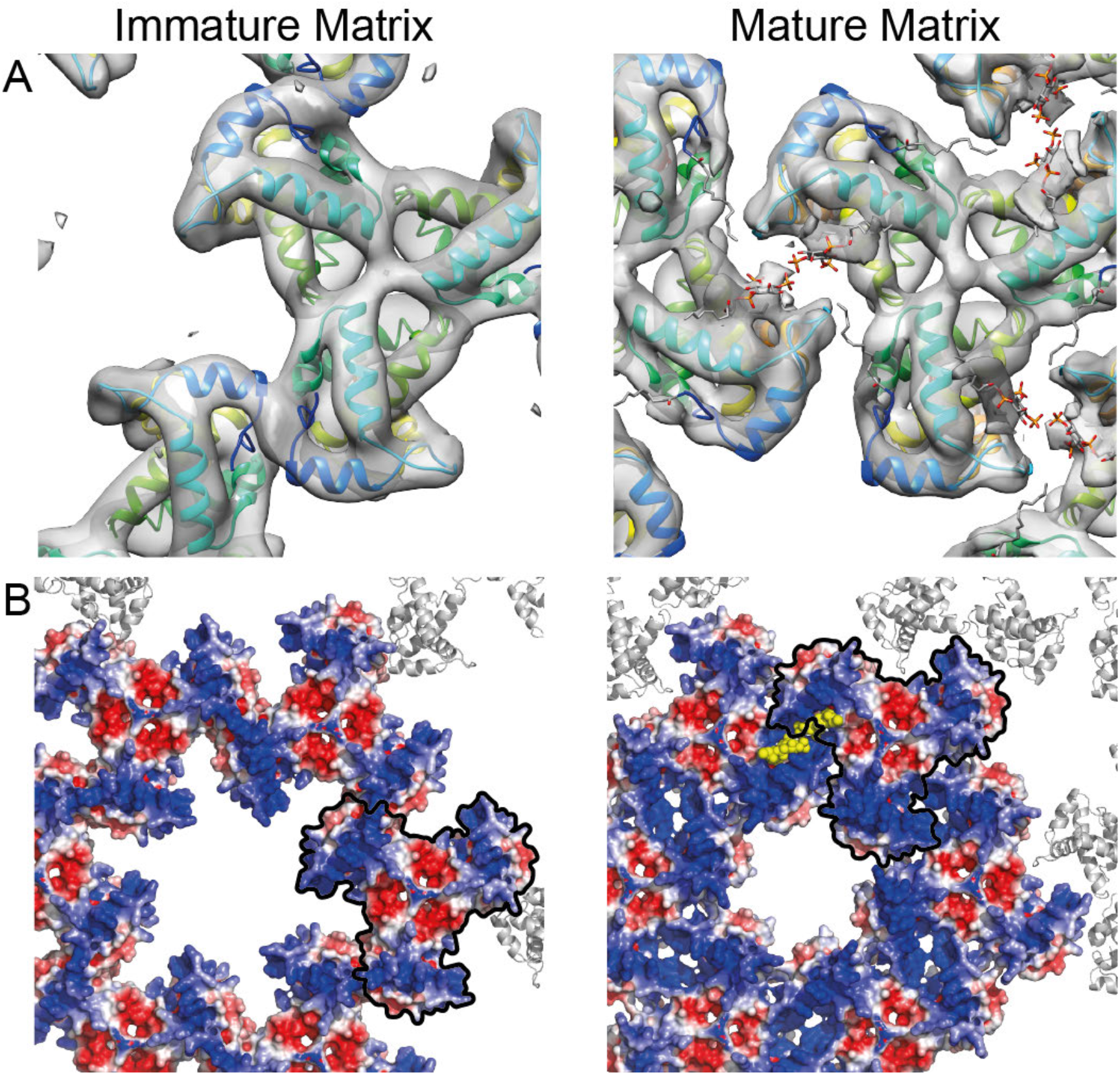
Comparison of the immature and mature HIV-1 MA lattices. **(A)** Immature and mature MA lattices are shown with one trimer aligned. In the immature lattice, contact with neighbouring trimers is mediated by N-terminal regions and the PI(4,5)*P_2_* binding site is empty. In the mature lattice, contact with neighbouring trimers is mediated by the region surrounding the occupied PI(4,5)*P_2_* binding site. **(B)** Electrostatic surface potential maps of the hexameric lattice of immature and mature MA trimers. The red (−5 kT/e) and blue (+5 kT/e) colours represent negatively and positively charged electric potentials. One MA trimer aligned as in (A) is outlined in black. The potential of the surface of MA facing the holes at the hexamer positions in the lattice changes from positive to neutral/negative during maturation. The hole becomes smaller. PI(4,5)*P_2_* is shown in yellow.

Taken together, our results (summarized in **Fig. S7**) revealed that MA – similar to CA and NC – undergoes dramatic structural maturation to form very different lattices in immature and mature HIV-1. Immature virions contain MA trimers packed together *via* their N-termini into a poorly-ordered hexagonal lattice containing basic-charged holes and disordered regions, which could accommodate the cytoplasmic tails of Env. The N-terminal myristate is inserted into the membrane. This immature MA structure provides a framework for understanding the established roles of MA in virus assembly and Env incorporation. Mature virions, in contrast, contain MA trimers packed together via their HBRs, resulting in a well-ordered hexagonal lattice with neutral holes. The maturation of MA is highly reminiscent of that of CA, which matures between two different hexameric protein lattices with two different functional roles (*3*). Upon maturation, both the protein arrangement and the lipid-binding properties of MA change. When bound to mature MA in the extended-lipid conformation, PI(4,5)*P_2_* is pulled outwards from the lipid bilayer (**Fig. S6B**) and one acyl chain is removed from the membrane to bind at the centre of the trimer-trimer interface. The energy barrier to the removal of acyl chains from the bilayer may be overcome by binding of PI(4,5)*P_2_* to MA and stabilization of the MA lattice. Our data imply that maturation of Gag results in the partial removal of ~2,500 of the ~150,000 lipids in the inner leaflet of the viral membrane (*20*).

Our data suggest that the structural maturation of Gag not only condenses the RNP and achieves assembly of a viral capsid core surrounding the genome, but also rearranges the membrane-bound MA layer and modulates the lipid bilayer itself. How these changes alter the properties of the virus remains to be elucidated, but we speculate that they impact not only MA and Env function, but also the physical properties of the viral envelope. The unexpected presence of a different well-defined MA structure in mature virions suggests that MA performs new roles in the mature virion or after entry into the target cell. We speculate that PI(4,5)*P_2_*-stabilized MA lattices may remain membrane associated after virion fusion and have a post-entry function, perhaps acting as signalling platforms to prepare the target cell for early HIV-1 replication.

## Materials and Methods

### Virus particle production and purification

HIV-1_NL4-3_ MA-SP1 PR+ particles were prepared by iodixanol gradient purification exactly as described in (*30*). cHIV and cHIV PR-particles were prepared for this study as follows: HEK 293T cells grown on 175-cm^2^ side-bottom tissue culture flasks were transfected with plasmid pcHIV (*18*) or pcHIV PR – (*33*) (70 μg of plasmid per flask) using a standard calcium phosphate transfection procedure. Culture media from producing cells were harvested at 48 h post-transfection and cleared by filtration through a 0.45-μm nitrocellulose filter. Particles from media were concentrated by ultracentrifugation through a 20% (wt/wt) sucrose cushion (90 min at 27,000 rpm in a Beckman SW32 rotor; Beckman Coulter Life Sciences). Pelleted particles were resuspended in PBS and stored in aliquots at −80°C.

### Cryo-electron tomography

Data imaging and processing were performed in parallel for cHIV PR- and cHIV particles. A suspension of purified viruses was diluted 10:1 with PBS solution containing 10-nm colloidal gold beads. An aliquot of 2.5 μl was applied on a glow-discharged C-Flat 2/2 3C grid (Protochips), and automatically back-side blotted and plunge-frozen into liquid ethane using the Leica EM GP 2. Grids were loaded into an FEI Titan Krios transmission electron microscope operated at 300 kV and imaging was performed on a Gatan BioQuantum K3 in super-resolution mode. Tomographic tilt series between −60° and +60° were collected using SerialEM-3.7.0 software (*34*) in a dose-symmetric scheme (*35*) with a 3° angular increment. The nominal magnification was 64 kx, giving a pixel size of 1.386 Å on the specimen. For other samples, data acquisition was performed similarly, on a Gatan Quantum 968 K2 in counting mode. All data acquisition parameters are listed in **Table S1**.

Frames were motion-corrected in IMOD-4.10.18 (*36*). K3 data were dose-filtered in IMOD-4.10.18, K2 data were dose-filtered in MATLAB (MathWorks). Exposure filtering was implemented according to the cumulative dose per tilt as described elsewhere (*37*). Fiducial-alignment of all tilt stacks was performed in IMOD-4.10.18 or older versions. Each tilt stack before dose-filtering was used to estimate the contrast transfer function (CTF) by CTFFIND-4.1.5 (*38*) or ctfplotter (*39*). Motion-corrected and dose-filtered tilt stacks were CTF-corrected by CTF multiplication and tomograms were reconstructed by weighted back-projected in NovaCTF-1.0.0 (*40*). The replicate of HIV-1_NL4-3_ PR-was CTF-corrected by phase flipping and reconstructed in IMOD-4.9.0.

### Subtomogram averaging

Subtomogram alignment and averaging were implemented using MATLAB (MathWorks) scripts derived from the TOM1 (*41*) and AV3 (*42*) packages as described previously (*30*). The missing wedge was modelled as the summed amplitude spectrum of background subtomograms for each tomogram, and was applied during alignment and averaging.

#### Structure determination from immature particles and cleavage mutants

Complete virus particles with distinct Gag layers were identified in 8-fold binned (bin8) tomograms and the centre of each virus was manually marked using “Volume Tracer” in UCSF Chimera (*43*). The irregular packing, small size, and proximity of MA to both CA and the membrane, make it challenging to solve the MA structure. To reduce the required search space, we first determined the structure of the CA layer as previously described (*19*) and thereby determined the location of CA hexamers. These positions were used to derive initial estimates of the position and orientation of MA.

To derive CA positions for HIV-1_NL4-3_ PR-, cHIV MA-SP1 and cHIV MA-CA, we made used of CA alignments from the published datasets (*19, 30*), performing minor reprocessing as necessary to ensure that as many CA positions as possible were retained during alignment.

To derive CA positions for cHIV PR-, HIV-1_NL4-3_ MA-SP1, and for a replicate of HIV-1_NL4-3_ PR-, the radius of the CA shell was measured and a mesh of initial extraction points was seeded on a spherical surface at this radius using MATLAB or a custom plug-in “Pick Particle” for UCSF Chimera (https://www2.mrc-lmb.cam.ac.uk/groups/briggs/resources/). An initial average of subtomograms from one bin8 tomogram was used as a starting reference for three iterations of translational and rotational alignment with a 50-Å low-pass filter and no symmetry applied. A reference-free structure was obtained and two further iterations were performed with 6-fold symmetry. All CA subtomograms were aligned against the 6-fold symmetrized reference at bin8 and further aligned at bin4 with 50-Å and 40-Å low-pass filters, respectively. Duplicated positions in the CA lattice were removed and then the lattice completeness was inspected manually. CA hexamers that were mis-aligned or with low CCC values were cleaned manually and then the dataset was split by virus particle number into two halves for further independent processing at bin2 and bin1, adapting the low-pass filter and angular search range appropriately while monitoring the resolution reported from the Fourier shell correlation (FSC).

We then extracted subtomograms at the expected radial position of MA, at locations directly above the 3-fold symmetry axes of the CA lattice, from bin2 tomograms. Preliminary orientations were assigned based on the orientation of the CA 3-fold symmetry axis. An initial reference was obtained by directly averaging the extracted subtomograms. For HIV-1_NL4-3_ PR-, alignments were tested applying C1 or C3 symmetry, both of which showed clear 3-fold symmetry. We therefore proceeded for all samples with alignments starting with 25 Å low-pass filter and C3 symmetry. After each iteration of alignment, we calculated the pairwise distance of all trimers in the dataset, and plotted the relative positions of all neighbouring MA trimers within about 8.6 nm distance (the typical MA trimer-trimer distances we observed were 5.6 nm for immature-like and 5.1 nm for mature-like particles) (**Fig. S1**). We averaged MA trimers for which all three neighbouring MA trimers were also present in the dataset to generate a new reference for the next alignment. After three further iterations of alignment at bin2, we again selected the MA trimers for which all three neighbouring MA trimers were present, and aligned only this subset at bin1 until the resolution no longer improved.

Resolution was measured from the phase randomization-corrected FSC between the two independent half datasets at 0.143 cut-off, using a mask considering only the central MA trimer. For structures at > 8 Å resolution, the combined maps were filtered and sharpened by an empirically determined B factor. For structures at < 8 Å resolution, the combined maps were locally filtered and sharpened by using a B factor estimated by the Guinier plot in Relion (*44*).

#### Structure determination from mature particles

In mature virions, CA cannot be used to derive approximate MA positions. We therefore segmented viral membranes using ilastik-1.3.3 (*45*) and extracted subtomograms at the inner surface of the membrane in bin6 tomograms. These were averaged and an arbitrary round density was placed at the centre of the box to facilitate initial convergence. Iterative alignments with a 50 Å low-pass filter was performed converging to a regular hexameric lattice. Bin4 subtomograms were then extracted and aligned against the 6-fold symmetrized reference. Duplicated positions in the MA lattice were removed and the dataset was split by virus particle number into two halves for further independent processing. Based on the positions of the aligned hexamers, the coordinates and orientations of trimers were calculated, and subtomograms containing trimers were extracted from bin2 data. Further alignments at bin2 and bin1 were performed until the structure converged. A biological replicate was performed using the 50 Å filtered reference which converged to the same structure, and the two datasets were therefore combined from bin2 to generate the final structure.

### Structure interpretation

The crystal structure (1HIW) of an MA trimer (*5*) was fitted into the immature and mature MA structures as a rigid body. To analyse the Myristate and PI(4,5)*P_2_* binding sites, we fitted the NMR structures of MA monomers with Myristate either exposed (2H3Q) or sequestered (2H3I) (*12*).

The lattice maps of CA hexamers and MA trimers were generated using the PlaceObject plugin (*46*) for UCSF Chimera. The electrostatic surfaces of immature and mature MA lattices were calculated in the APBS plugins (*47*) for PyMOL (The PyMOL Molecular Graphics System, Version 1.8 Schrödinger, LLC.).

## Acknowledgments

We thank Wim Hagen, Dustin Morado, Aaron Tan, Florian Schur, and Simone Mattei for technical assistance, and Petr Chlanda and Leo James for helpful discussions. This study made use of electron microscopes at EMBL and the MRC-LMB EM Facility, as well as high-performance computing resources set-up and maintained by EMBL IT Services and LMB Scientific Computing; we thank the staff who maintain those resources

## Author contributions

KQ, BM, HGK and JAGB conceived the project. KQ, ZK, VZ, BM, HGK and JAGB designed experiments. VZ, MAÖ and BG purified virus particles. KQ and ZK performed electron microscopy. KQ and ZK performed computational image processing. KQ and JAGB wrote the original draft with assistance from BM and HGK which was edited and reviewed by all authors. KQ prepared figures. BM, HGK and JAGB obtained funding, provided supervision and managed the project

## Data and materials availability

The immature and mature lattice structures and representative tomograms are deposited in the Electron Microscopy Data Bank (EMDB) under accession codes EMD-XXXXX - EMD-XXXXX. The associated molecular models are deposited in the protein data bank (PDB) under accession XXXX and XXXX.

---

**Fig. S1.**
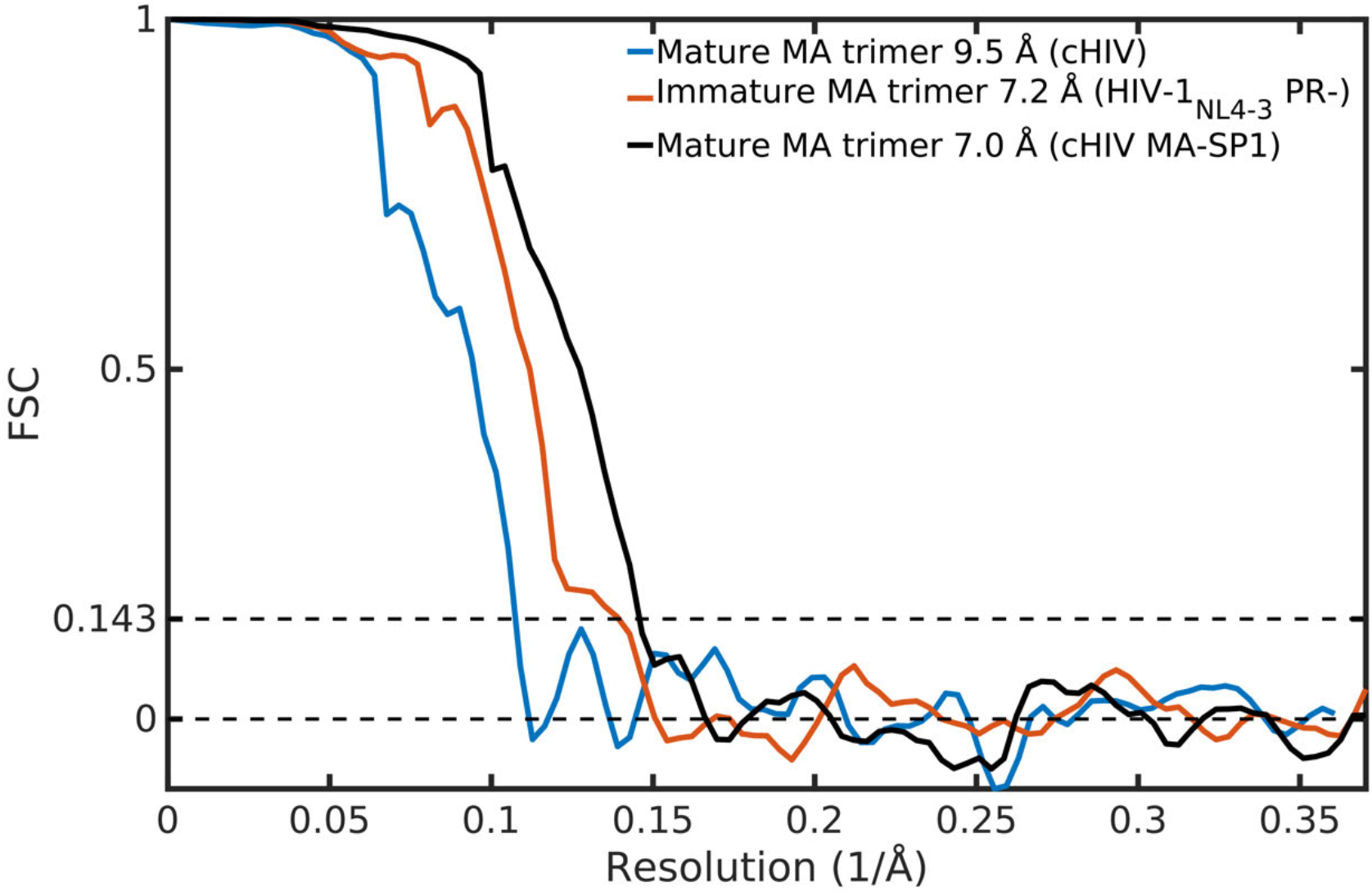
Fourier shell correlation (FSC) curves to measure the resolution of MA lattice structures. Phase randomization-corrected FSC curves were calculated between two independent halfmaps masked by a cylinder including the central MA trimer. Resolution values were reported at the 0.143 threshold. cHIV (mature, blue), HIV-1_NL4-3_ PR- (immature, red) and cHIV MA-SP1 (mature, black).

**Fig. S2.**
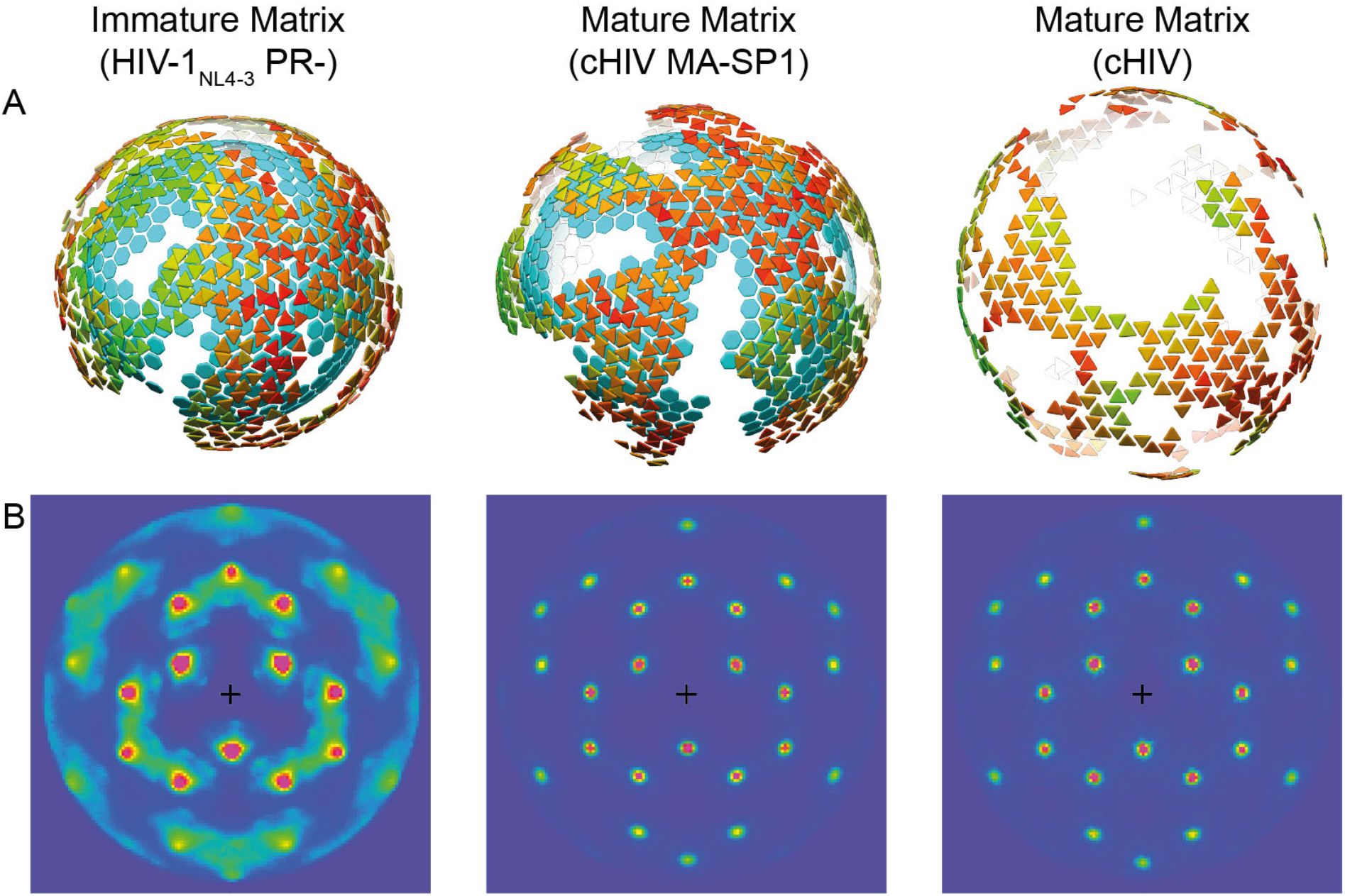
Comparison of immature and mature HIV-1 MA lattices. **(A)** Further examples of lattice maps derived from subtomogram averaging, for the immature MA lattice (left), for the mature MA lattice from MA-SP1 (center), and for the mature MA lattice derived from mature cHIV particles (right). Lattice maps are displayed as in Fig. 2G and Fig. 3F, Positions and orientations of MA trimers are shown as triangles colored on a scale from red (lower cross correlation to average structure) to green (higher cross correlation to average structure). Where present, the positions of immature CA hexamers are indicated as blue hexagons to illustrate the relationship between MA and CA layers. **(B)** Heatmaps of neighbour plots for immature and mature MA trimers to illustrate the regularity of the lattices in (A). Peaks (red) denote the positions of all neighbouring MA trimers relative to a central trimer (position marked by a black cross), averaged for all trimers in the lattice. Peaks representing the first, second and third order neighbours are distinctly visible for both immature and mature lattices. Long-distance order is only observed in the mature MA lattice. Sharper peaks correspond to a more regular lattice.

**Fig. S3.**
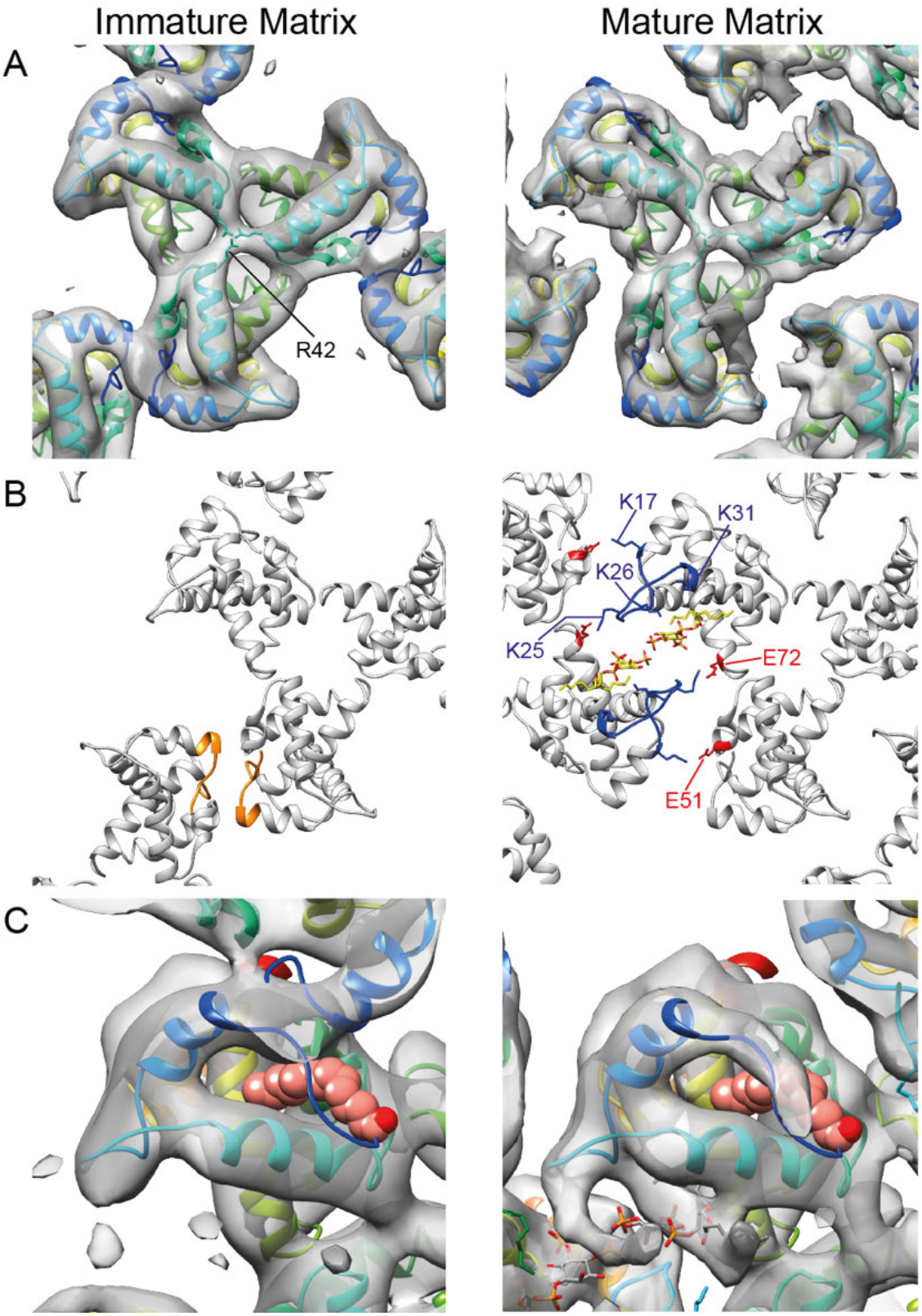
Further comparison of immature and mature HIV-1 MA lattices. **(A)** Immature and mature (MA-SP1) MA lattices are shown with one trimer aligned. In both cases the central pore of the MA trimer is occluded, and is surrounded by a positively charged electric potential contributed by three arginines (R42). This site corresponds to that previously observed to interact with phosphatidylserine (PS), phosphatidylethanolamine (PE) and phosphatidylcholine (PC), but the available space after trimerization cannot accommodate a lipid. **(B)** Immature and mature MA shown in cartoon representation. Left: in the immature MA lattice, inter-trimer interactions are mediated by N-terminal residues that change conformation between myristate-exposed and myristate-inserted states (residues 1-11, orange). Right: in the mature MA, basic residues in the HBR loop (blue) face acidic residues in the N-terminus of helix 4 (72E) and the 3_10_ helix (51E) of the adjacent MA monomer (red). PI(4,5)*P_2_* is shown in yellow. **(C)** An enlargement of MA monomer from the immature (left) and mature (right) MA trimers, shown as grey isosurface, and fitted with the NMR structure (2H3I) of MA with sequestered myristate. The myristate is shown as pink spheres. No significant EM density was observed in the myristoyl pocket, indicating that the myristate is not sequestered.

**Fig. S4.**
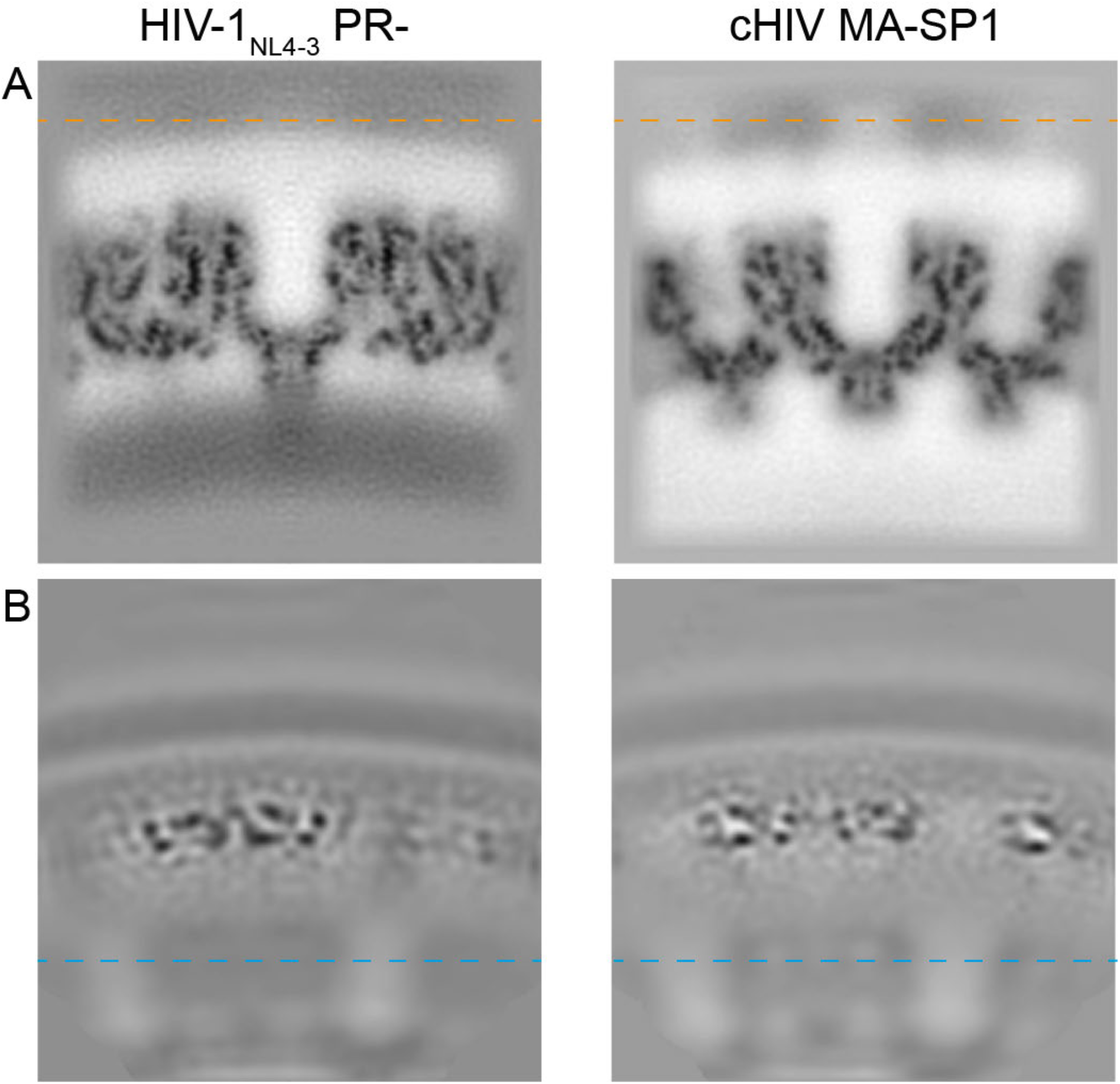
Orthoslices perpendicular to the viral membrane through different reconstructions show no fixed relationship between MA and CA layers and no ordered density is found between CA and MA layers. **(A)** Alignments focus on the CA lattice resulting in blurred MA layers (MA layer at position of orange dashed line). Published CA structures are shown on the left (HIV-1_NL4-3_ PR-, EMD-4017) and right (cHIV MA-SP1, EMD-0164). **(B)** From the same datasets, with alignments focussed on the MA layer, the CA layer becomes blurred (CA position at cyan dashed line).

**Fig. S5.**
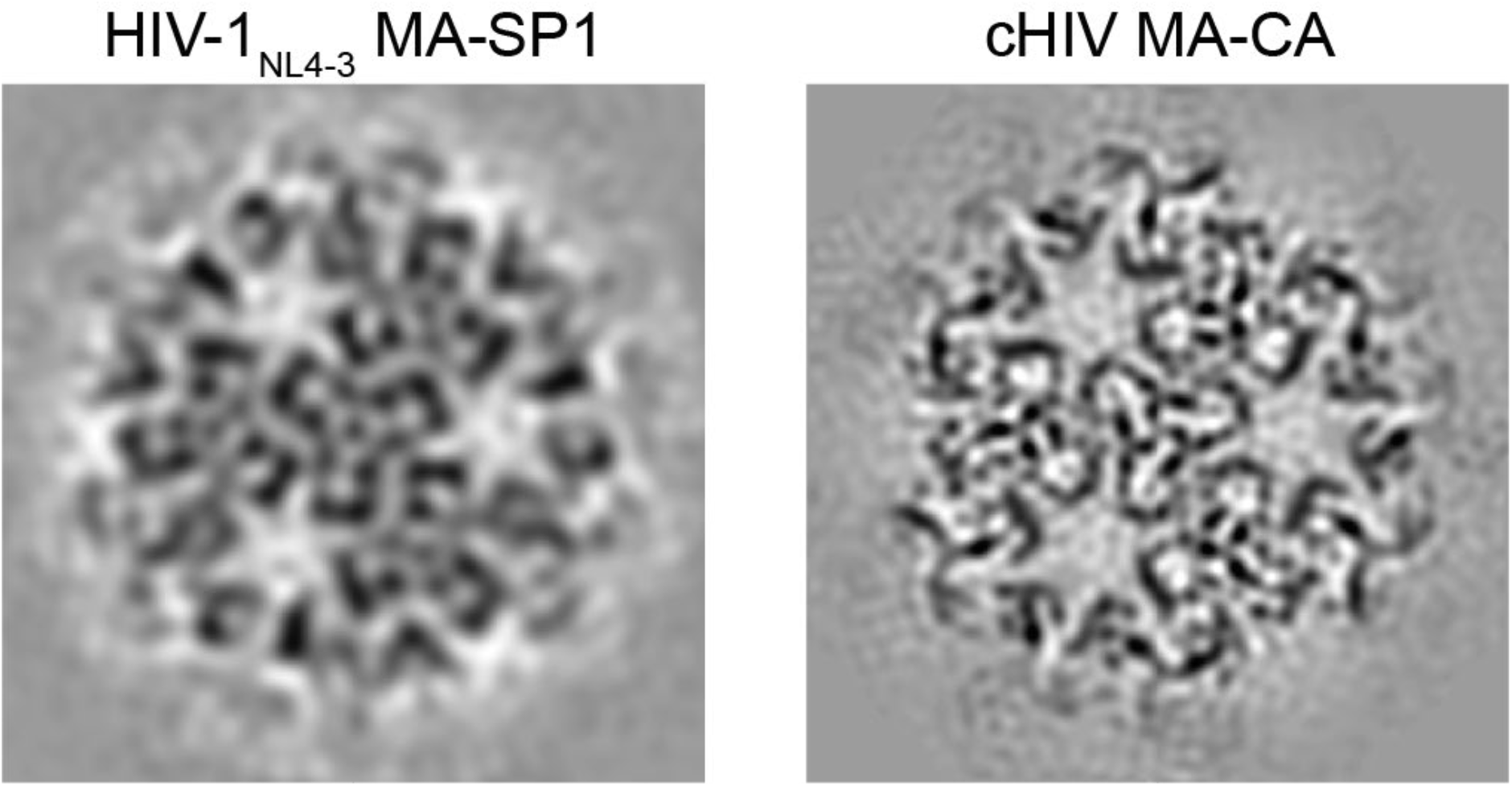
Mature HIV-1 MA structures in HIV-1_NL4-3_ MA-SP1 (left) and cHIV MA-CA (right), derived as additional control experiments. As expected, orthoslices through the MA layer show an identical mature MA arrangement to that seen in lattices in mature cHIV and cHIV MA-SP1 particles as shown in Figure 3A.

**Fig. S6.**
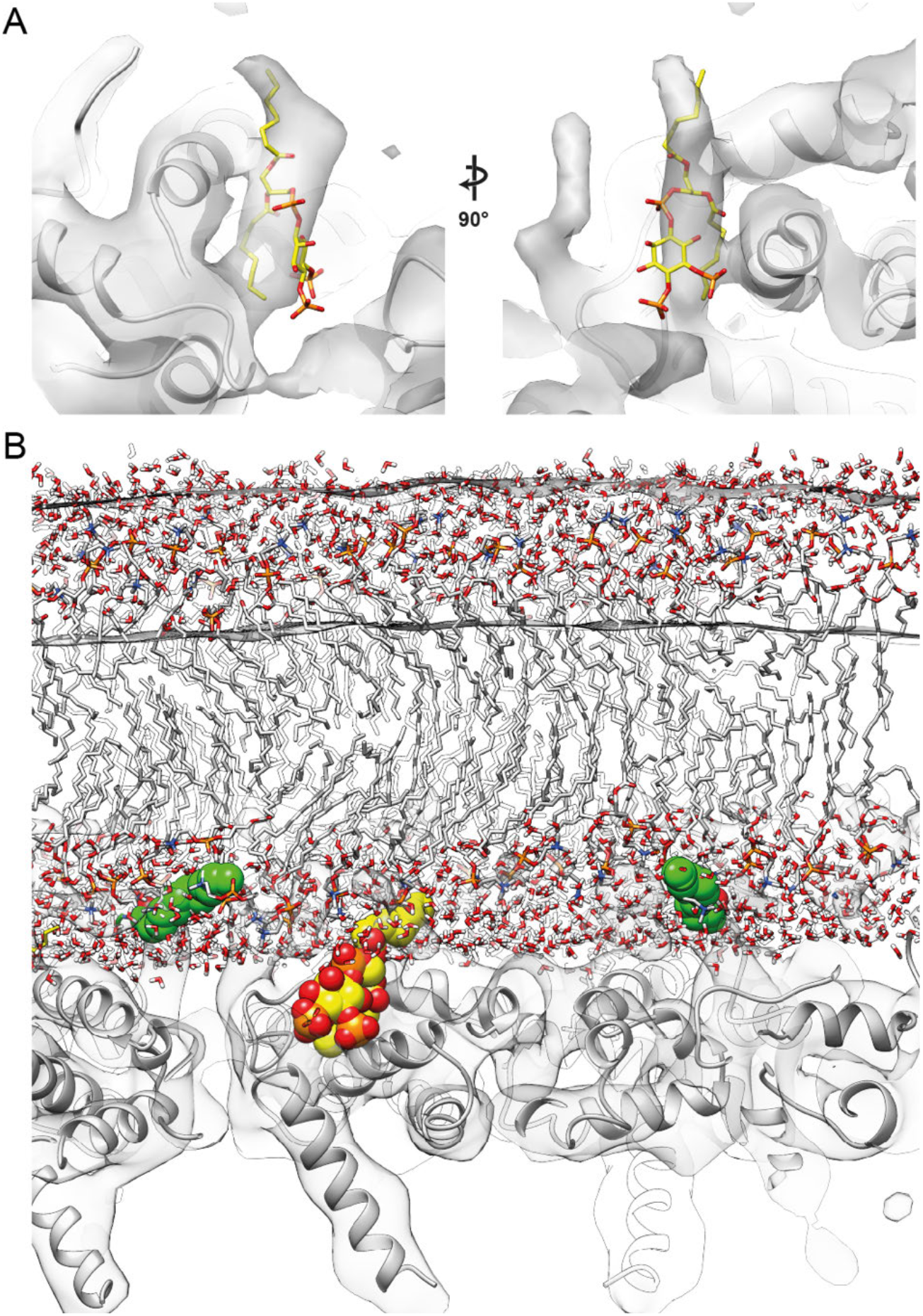
The position of bound PI(4,5)*P_2_* in the mature HIV-1 MA lattice. **(A)** PI(4,5)*P_2_* (yellow and red) from the NMR structure (PDBID: 2H3V) was fitted as a rigid body together with the MA model (PDBID: 2H3Q) into the mature MA density (transparent grey). The 1’ acyl chain points upwards towards the membrane while and the 2’ acyl chain (down) is sequestered in the MA layer. **(B)** The position of the mature HIV-1 MA lattice relative to the bilayer is shown by fitting a model for the bilayer (*48*) into the membrane density in the MA structure. This representation highlights that the headgroup of PI(4,5)*P_2_* (yellow and red spheres) is moved outwards from the bilayer and embedded in the binding pocket, the 1’ acyl chain (up) is partially pulled out from the bilayer and passes through the lipid headgroup layer, and the 2’ acyl chain is completely removed from the bilayer. The exposed N-terminal myristate (green spheres) is fully inserted into the inner leaflet of lipid bilayer.

**Fig. S7.**
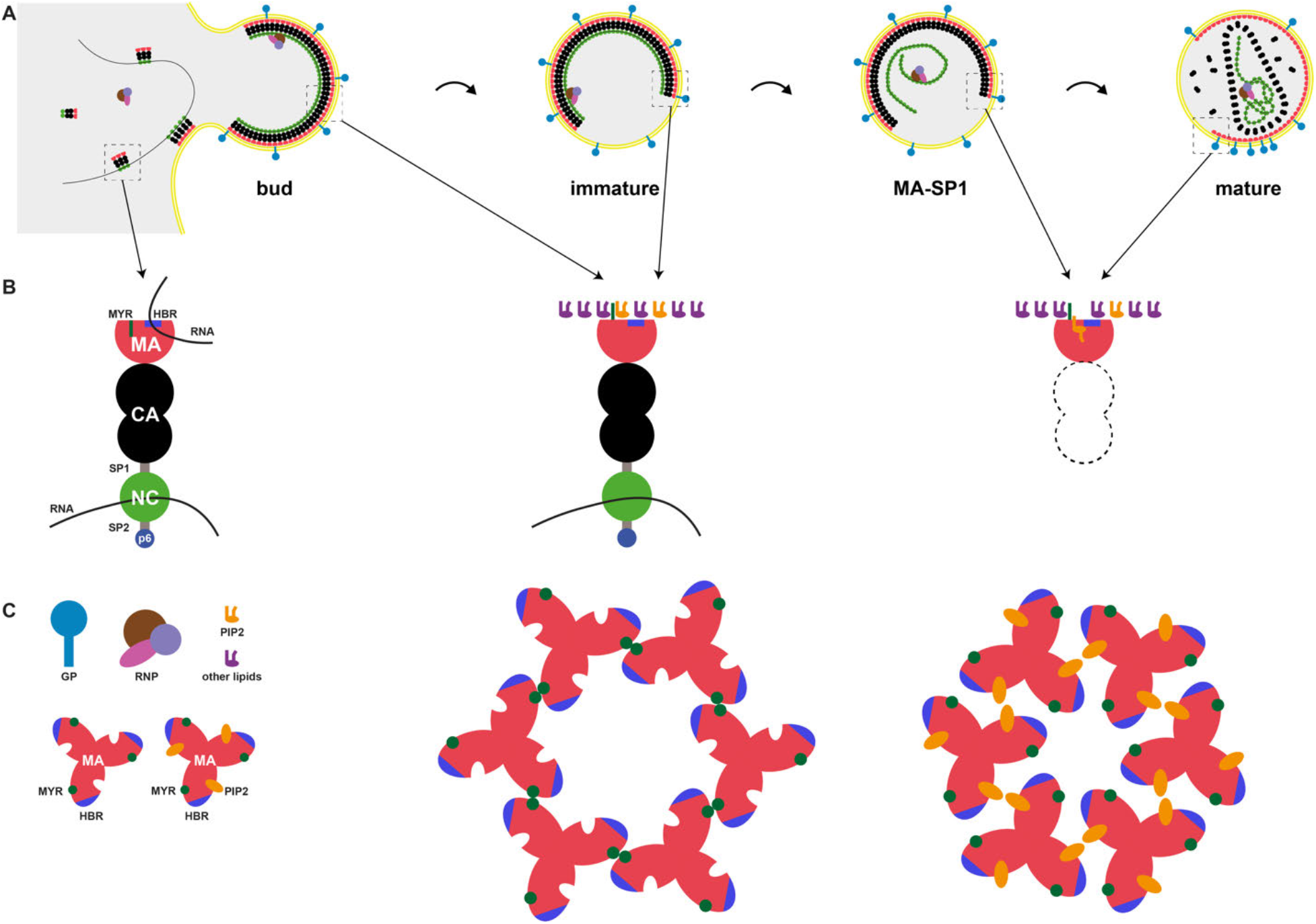
Schematic illustrations of MA-MA and MA-membrane interactions during assembly and maturation. **(A)** Gag in the cytosol binds RNA and assembles at the plasma membrane leading to formation of a bud. During or after virus release, the immature virus undergoes proteolytic maturation to form the mature virion. The MA-SP1 cleavage mutant may not represent a true maturation intermediate. Boxed regions are highlighted in (B). **(B)** Interactions between Gag and the plasma membrane via PI(4,5)*P_2_* (PIP2) and myristate (MYR). In the cytosol, the N-terminal myristate is sequestered in MA and RNA is bound to the HBR. In the assembling and immature virion, RNA is released from the HBR while the myristate is exposed and inserted into the plasma membrane. After proteolytic cleavage between SP1 and NC, the PI(4,5)*P_2_* head group and unsaturated 2’ acyl chain are pulled out from the PM and sequestered in a positively charged pocket on MA. **(C)** Structural maturation of HIV-1 MA. In the immature MA lattice, MA trimer-trimer interactions are formed by the N-terminal domain in the vicinity of the exposed myristate moiety, while the PI(4,5)*P_2_* binding pocket is empty. In the mature MA lattice, PI(4,5)*P_2_* is bound to its binding pocket at the side of MA and MA trimer-trimer interactions are formed by the HBR and PI(4,5)*P_2_*.

**Table S1.** Cryo-EM data acquisition and image processing.

| Sample | cHIV<br>PR-Transfec<br>tion I | cHIV<br>Transfec<br>tion I | cHIV<br>Transfec<br>tion II | HIV-1<sub>NL4-3</sub><br>MA-SP1 | HIV-1<sub>NL4-3</sub><br>PR-Replicate | HIV-1<sub>NL4-3</sub><br>PR-<sup>a</sup> | cHIV<br>MA-SP1<sup>b</sup> | cHIV<br>MA-CA<sup>b</sup> |
| --- | --- | --- | --- | --- | --- | --- | --- | --- |
| **Data acquisition** |  |  |  |  |  |  |  |  |
| Microscope | FEI Titan Krios | FEI Titan Krios | FEI Titan Krios | FEI Titan Krios | FEI Titan Krios | FEI Titan Krios | FEI Titan Krios | FEI Titan Krios |
| Voltage (keV) | 300 | 300 | 300 | 300 | 300 | 300 | 300 | 300 |
| Energy-filter (eV) | 20 | 20 | 20 | 20 | 20 | 20 | 20 | 20 |
| Detector | Gatan BioQuantum K3 | Gatan BioQuantum K3 | Gatan BioQuantum K3 | Gatan K2 Summit | Gatan K2 Summit | Gatan K2 Summit | Gatan K2 Summit | Gatan K2 Summit |
| Pixel size (Å) | 1.386 | 1.386 | 1.386 | 1.379 | 1.35 | 1.35 | 1.35 | 1.35 |
| Defocus range (microns) | -2.0 to -4.5 | -4.5 to -7.5 | -4.5 to -7.5 | -1.1 to -3.1 | -2.0 to -4.0 | -1.5 to -5.0 | -1.5 to -5.0 | -1.5 to -5.0 |
| Acquisition scheme | -60/60°, 3° | -60/60°, 3° | -60/60°, 3° | -60/60°, 3° | -60/60°, 3° | -60/60°, 3° | -60/60°, 3° | -60/60°, 3° |
| Total Dose (electrons/Å<sup>2</sup>) | ~120 | ~120 | ~120 | ~120 | ~95 | ~120-221 | ~145 | ~150 |
| Dose rate (electrons/Å<sup>2</sup>/sec) | ~13.0 | ~13.0 | ~13.0 | ~2.1 | ~4.6 | ~1.5 – 5.5 | ~2.3 | ~2.3 |
| Frame number | 10 | 10 | 10 | 10 | 5 | 10-12 | 10 | 21 |
| Tomogram number | 53 | 25 | 20 | 26 | 32 | 74 | 65 | 70 |
| **Image processing** |  |  |  |  |  |  |  |  |
| Virus particles | 278 | 122 |  | 106 | 155 | 476 | 350 | 302 |
| Subtomogram (set A/B) | 7,254/6,769 | 15,384/14,895 |  | 12,372/11,756 | 24,663/24,370 | 10,980/11,437 | 9,601/9,985 | 14,150/15,333 |
| Symmetry | C3 | C3 |  | C3 | C3 | C3 | C3 | C3 |
| Resolution at 0.143 FSC (Å) | 13.3 | 9.5 |  | 10.6 | 9.6 | 7.2 | 7.0 | 6.6 |
| B factor | -500 | -500 |  | -500 | -500 | -597 | -521 | -511 |

<sup>a</sup> Data published previously is reinvestigated (19).
<sup>b</sup> Data published previously is reinvestigated (30).

